# Reconstruction of cell spatial organization based on ligand-receptor mediated self-assembly

**DOI:** 10.1101/2020.02.13.948521

**Authors:** Xianwen Ren, Guojie Zhong, Qiming Zhang, Lei Zhang, Yujie Sun, Zemin Zhang

## Abstract

Single-cell RNA sequencing (scRNA-seq) has revolutionized transcriptomic studies by providing unprecedented cellular and molecular throughputs, but spatial information of individual cells is lost during tissue dissociation. While imaging-based technologies such as *in situ* sequencing show great promise, technical difficulties currently limit their wide usage. Since cellular spatial organization is inherently encoded by cell identity and can be reconstructed, at least in part, by ligand-receptor interactions, here we present CSOmap, a computational strategy to infer cellular interaction from scRNA-seq. We show that CSOmap can successfully recapitulate the spatial organization of tumor microenvironments for multiple cancers and reveal molecular determinants of cellular interactions. Further, CSOmap readily simulates perturbation of genes or cell types to gain novel biological insights, especially into how immune cells interact in the tumor microenvironment. CSOmap can be widely applicable to interrogate cellular organizations based on scRNA-seq data for various tissues in diverse systems.

## Main Text

High-throughput single-cell RNA sequencing has emerged as a revolutionary approach to dissect cellular compositions and characterize molecular properties of complex tissues^1^, and has been applied to a wide range of fields resulting in profound discoveries^2^. However, spatial information of individual cells is lost during the process of tissue dissociation. While it is paramount to investigate the molecular composition of individual cells in the spatial contexture, current methods such as RNA hybridization^3^, *in situ* sequencing^4^, immunohistochemistry^5^, and purifying predefined subpopulations for subsequent transcriptomic profiling^6^ are limited by the throughput and complex experimental procedures that are only accessible by a handful of laboratories. The combination of scRNA-seq with *in situ* RNA patterns or tissue shapes provides computational solutions for high-throughput mapping the spatial locations of many individual cells^7–10^, but this type of methods relies on the availability of such spatial references, limiting their wide applications.

The spatial organization of individual cells has recently been shown to be self-assembled via ligand-receptor interactions^11, 12^, implying that cellular spatial organization are inherently encoded by their identity. Thus, we argue that the spatial relationship of cells may be *de novo* reconstructed, at least in part, by integrating scRNA-seq data with known ligand-receptor interaction information. We hypothesize that: 1) the potential of cellular interactions can be approximated by the abundance of interacting ligands and receptors and their affinity; 2) cells with high interacting potentials tend to locate in close proximity; 3) cells compete for their interacting partners due to physiological and spatial constraints. We formulate these hypotheses as a mathematical optimization model (named as CSOmap) that predicts coordinates of each cell in a three-dimensional pseudo-space based on input scRNA-seq data and known ligand-receptor interactions^13, 14^ (Figure 1).

**Figure 1.**
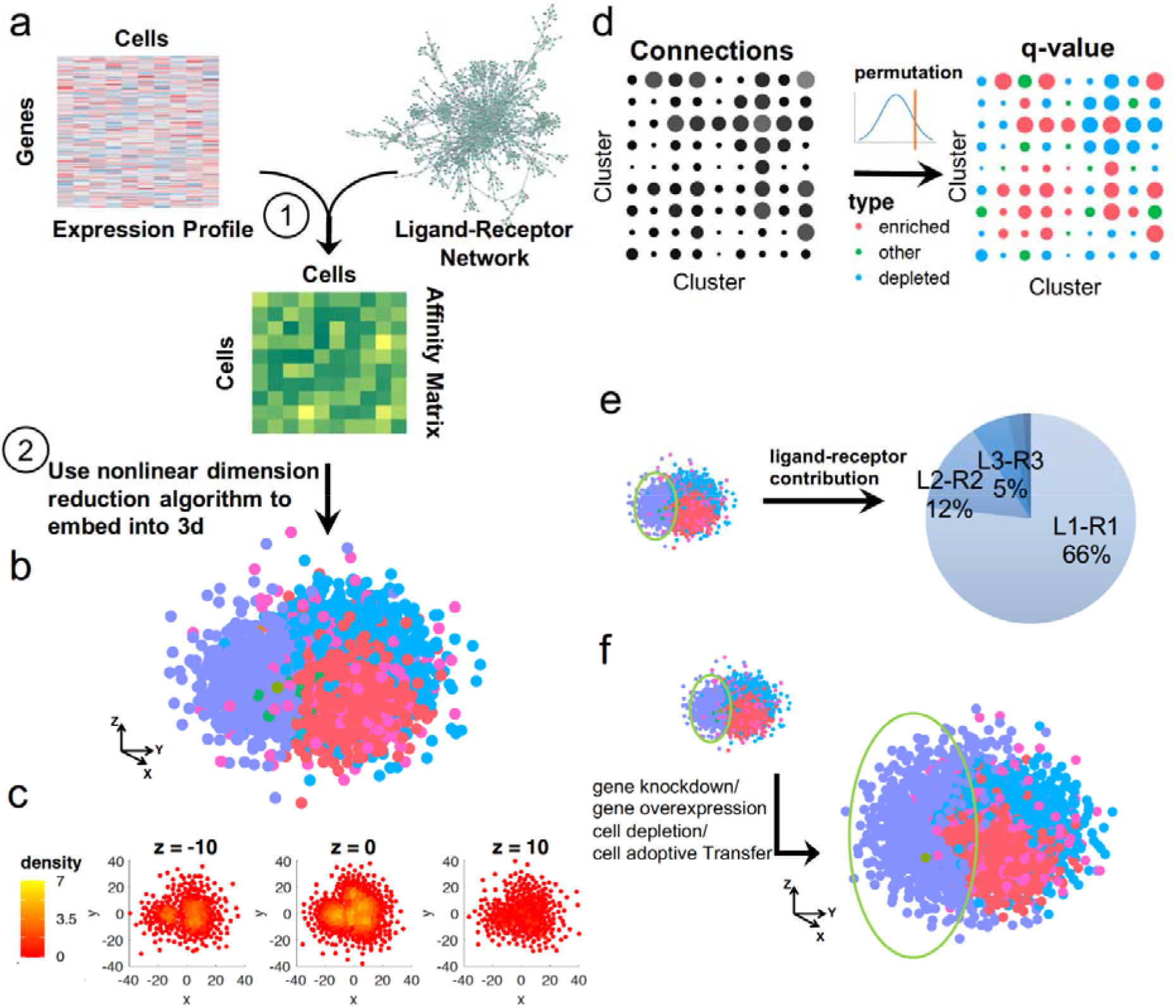
Schematics of single-cell spatial reconstruction by CSOmap. (a) CSOmap takes the gene-by-cell expression matrix generated by scRNA sequencing and the known ligand-receptor network as inputs, upon which a cell-by-cell affinity matrix is estimated. (b) The inherently high-dimensional cell-by-cell affinity matrix is embedded into a three-dimensional space via resolving cell competitions. (c) Density can be estimated for individual cells based on their three-dimensional coordinates obtained from (b), which allows the identification of spatially-defined cell clusters. (d) Given the cell cluster labels, the number of connections among cell clusters and their statistical significance can be summarized and evaluated by CSOmap. (e) For a pair of cell clusters, the contributions of each ligand-receptor pair to their interactions can be calculated. (f) CSOmap allows *in silico* interference of the original dataset including gene knockdown/overexpression and cell depletion/adoptive transfer to examine the corresponding effects on cellular spatial organizations.

The algorithmic process is composed of two main steps. The first is to estimate the cellular interacting potentials by integrating thousands of ligand-receptor pairs, resulting in a cell-by-cell affinity matrix (Figure 1a). The second is to embed the inherently high-dimensional affinity matrix into three-dimensional space (Figure 1b). The limited availability of space determines that it is not feasible to position cells with the same interacting potentials equally close to their partner. Thus, we applied Student’s t-distribution to resolve the cell competition problem, enlightened by the widely used visualization technique t-SNE^15^. After embedding cells into three-dimensional space, spatial structures/patterns of cells can be analyzed by density-based clustering^16^ (Figure 1c), connections and corresponding statistical significance among predefined cell types can be summarized (Figure 1d), and the dominant ligand-receptor pairs underlying a specified pair of cell types can be calculated (Figure 1e). When a critical gene or cell population was determined, CSOmap can be further applied to examine the effects of *in silico* perturbations including gene overexpression or knock-down and cellular depletion or adoptive transfer (Figure 1f).

We first evaluated CSOmap on various publicly available scRNA-seq datasets. In a scRNA-seq dataset of pancreas^17^, the spatial separation of the endocrine and exocrine compartments provides a natural reference for assessing the performance of CSOmap. On both the human and mouse pancreatic scRNA-seq data, CSOmap successfully recapitulated such spatial separation (Supplementary Fig. 1), with endocrine cells forming one structure and exocrine cells composing the other compartment. The visual separation was further supported by random permutation-based statistical testing. Then CSOmap also successfully reconstructed the early maternal-fetal interface based on scRNA-seq data of human placenta and decidua^18^, *i.e*., fetal placenta cells, rather than maternal blood cells, showing significant interactions with maternal decidua cells (Supplementary Fig. 2a). Quantitative evaluation based on the scRNA-seq data of mouse liver lobules^8^ showed that CSOmap reached high consistence (*R* = 0.85, *p* < 0.01, Spearman correlation, Supplementary Fig. 2b) with the reference^8^. Due to the inherent difficulty to dissociate endothelial cells from liver cells, paired-cell sequencing has been customized to resolve the spatial positions of endothelial cells within liver lobules^19^. *In silico* predictions by CSOmap reached consistent results with paired-cell sequencing (*R* = 0.73, *p* < 0.04, Spearman correlation, Supplementary Fig. 2c). Systematic evaluation based on the Tabula Muris datasets^20^ demonstrated that CSOmap could reproduce the organ-level separations by revealing significantly higher intra-organ cellular interactions than inter-organ interactions for almost all organs except tongue (15/16, 93.75%, Supplementary Fig. 3), of which 176 out of 199 interacting cell type pairs were from different cell types. Such successful applications clearly show the effectiveness, robustness, and wide applicability of CSOmap for multiple organs from different organisms and different technical platforms.

We further experimentally validated the performance of CSOmap. First, we dissected a liver tumor sample into tumor edges and cores and applied scRNA-seq separately. With the scRNA-seq data only, CSOmap successfully reconstructed a cell spatial organization with tumor core cells locating in the center and tumor edge cells outside (Figure 2a). The spatial separation was statistically significant by quantitatively evaluating the distance of tumor core and edge cells to the center of the pseudo-space reconstructed by CSOmap (Figure 2b). The spatial patterns of genes encoding heat-shock proteins (Hsps) (Hsp40, Hsp70 and Hsp90) revealed by CSOmap, *i.e.*, with high expression level in the center while low expression level near the edge (Figure 2c), were further experimentally confirmed by immunohistochemical (IHC) staining on independent liver tumor samples (Figure 2d and 2e), suggesting the effectiveness and robustness of CSOmap in deriving new biological insights.

**Figure 2.**
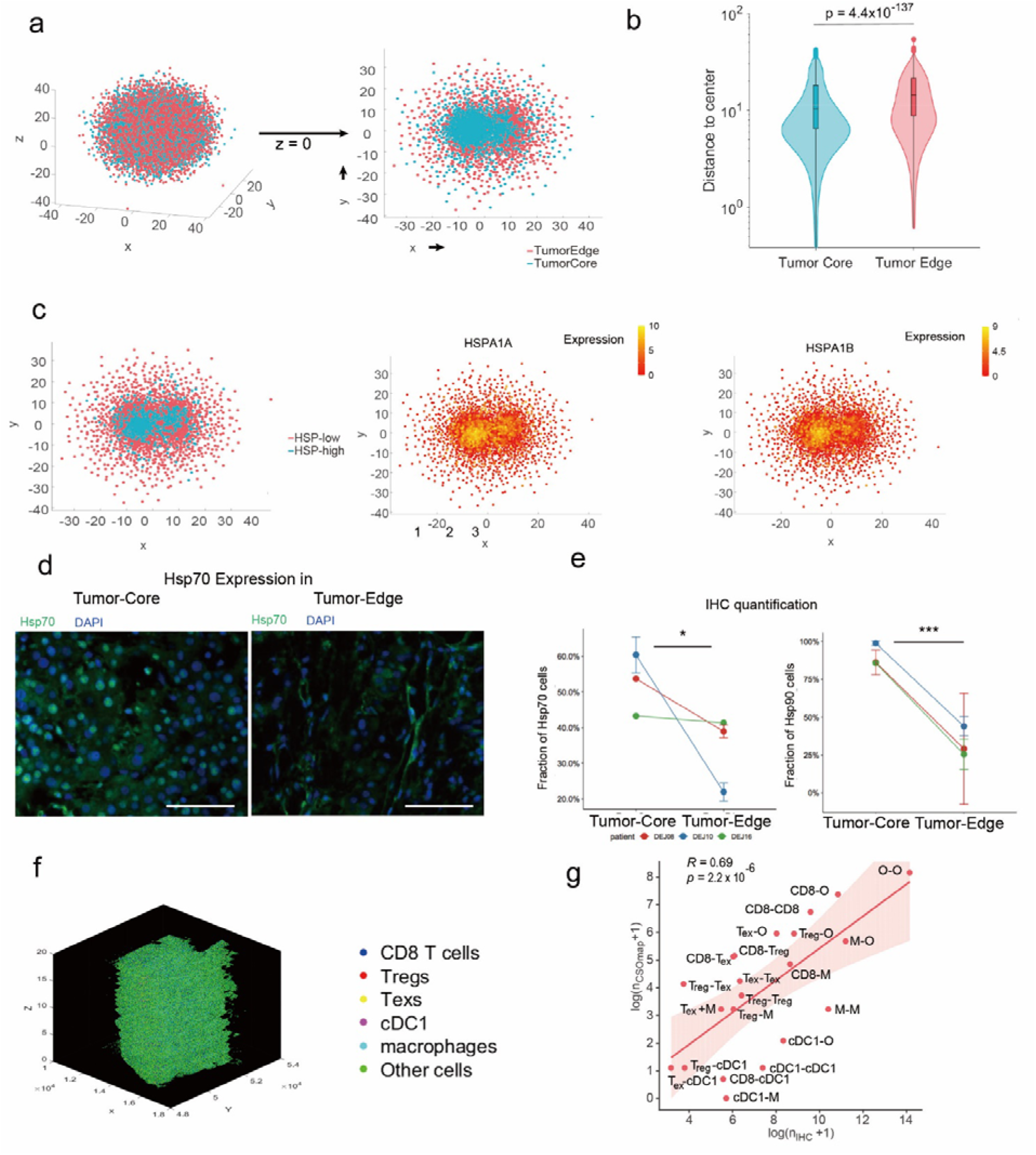
Performance of CSOmap in reconstructing the spatial organization of a liver tumor sample. (a) and (b) Tumor core cells tend to locate in the center of the pseudo-space reconstructed by CSOmap. (c) The CSOmap reconstruction revealed that genes encoding HSPs show spatial preference. (d) IHC staining on independent liver tumor samples confirmed the spatial preference of Hsp70. The scale bar represents 50 μm. (e) Quantification based on IHC images confirmed the statistical significance of the spatial preference of Hsp70 and Hsp90 (Student’s t-test, right tailed, *: p<0.05; ***: p < 0.01). (f) 3D plot of the tumor sample by stacking 19 IHC images together after manual rotation, in which six major cell types were discriminated by the corresponding markers. (g) Spearman correlation between cell connections based on IHC images (X-axis) and the CSOmap prediction (Y-axis). Treg: regulatory T cells (Foxp3+); Tex: exhausted T cell (PD-1+); CD8: CD8+PD-1-T cells; cDC1: CLEC9A+ dendritic cells; M: macrophages (CD68+); O: other cells. The median distance of the 3^rd^ nearest neighbor of all cells was used as the cutoff to determine whether two cells were spatially connected or not.

We then assessed the CSOmap results in a quantitative manner by simultaneously generating scRNA-seq and IHC staining data for a tumor sample derived from hepatocellular carcinoma. In brief, six cell types, including regulatory T cells (Tregs, marked by Foxp3+), exhausted CD8+ T cells (Texs, marked by PD-1+), CD8+PD-1^-^ T cells, type 1 dendritic cells (cDC1, marked by CLEC9A+), macrophages (marked by CD68+), and other cells, were labeled by specific antibodies on a 1cm × 1cm × 100um tumor tissue. This tumor tissue was consecutively spliced into 1cm × 1cm × 5um pieces for IHC staining and then the cell types and positions of 1,181,790 cells were recorded to serve as the reference for evaluating the performance of CSOmap (Figure 2f, and Supplementary Fig. 4). With 1,329 scRNA-seq profiles based on SMART-seq2^21^, CSOmap reached high concordance with the results of IHC analysis (Figure 2g, *R* = 0.69, *p* = 2.2 × 10-6, Spearman correlation) and recapitulated multiple cell-cell interactions exemplified by CD8 T cells-macrophages and Tregs-Texs pairing (Supplementary Fig. 5). After removing the potential biases introduced by the uneven cell counts of different cell types, the consistence score between IHC results and the CSOmap prediction was still 0.54 (*p* = 2.0 × 10^-4^, Spearman correlation) while the correlations based on random coordinate assignment and random gene pairs were −0.12 and 0.34, respectively.

Besides the spatial reconstruction, CSOmap can also provide important insights into the underlying molecular mechanisms. We applied CSOmap separately to a head and neck cancer (HNC) scRNA-seq dataset^22^ and a melanoma scRNA-seq dataset^23^. Based on the IHC images of the original report^22^, the spatial characteristics of HNC tumor microenvironment can be summarized as 1) malignant cells not subject to partial epithelial to mesenchymal transition (p-EMT) were located close to each other and formed a loose structure; 2) malignant cells subject to p-EMT were located at the interface between malignant cells and cancer-associated fibroblasts (CAFs); 3) CAFs were connected to each other and formed a compact structure (Figure 3a). CSOmap not only qualitatively recapitulates all these IHC characteristics (Figure 3b), but also highlighted the distinct spatial patterns of malignant cells between HNC and melanoma, *i.e.*, malignant cells in HNC tend to form a loose structure (adjusted *p* > 0.05, permutation-based test) while tumor cells in melanoma tend to form a compact structure (Figure 3c, adjusted p < 0.05, permutation-based test). By analyzing the dominant ligand-receptor pairs contributing to this spatial organization, we identified that the interactions between CD63 and TIMP1 contributed ~66% to the cellular interaction potential of melanoma malignant cells (Figure 3d) while HNC cells expressed CD63 and TIMP1 at much lower levels. Using CSOmap, we were able to readily perform “*in silico* perturbation” of CD63 and re-calculate the spatial characteristics. Indeed, *in silico* knocking down of CD63 expression levels in melanoma malignant cells resulted in the transition from compact to loose structures while overexpression of CD63 in HNC malignant cells resulted in compact structure (Figure 3e and 3f). The association of CD63 with the morphology of melanoma has been experimentally supported by a previous *in vivo* and *in vitro* study^24^, in which the mechanism underlying such association was attributed to the negative linkage between CD63 signaling and epithelial to mesenchymal transitions. This notion is recapitulated by CSOmap since the p-EMT program was observed in the HNC dataset but absent in melanoma, suggesting the effectiveness of CSOmap in spatial reconstruction and the potential in revealing the underlying molecular mechanism.

**Figure 3.**
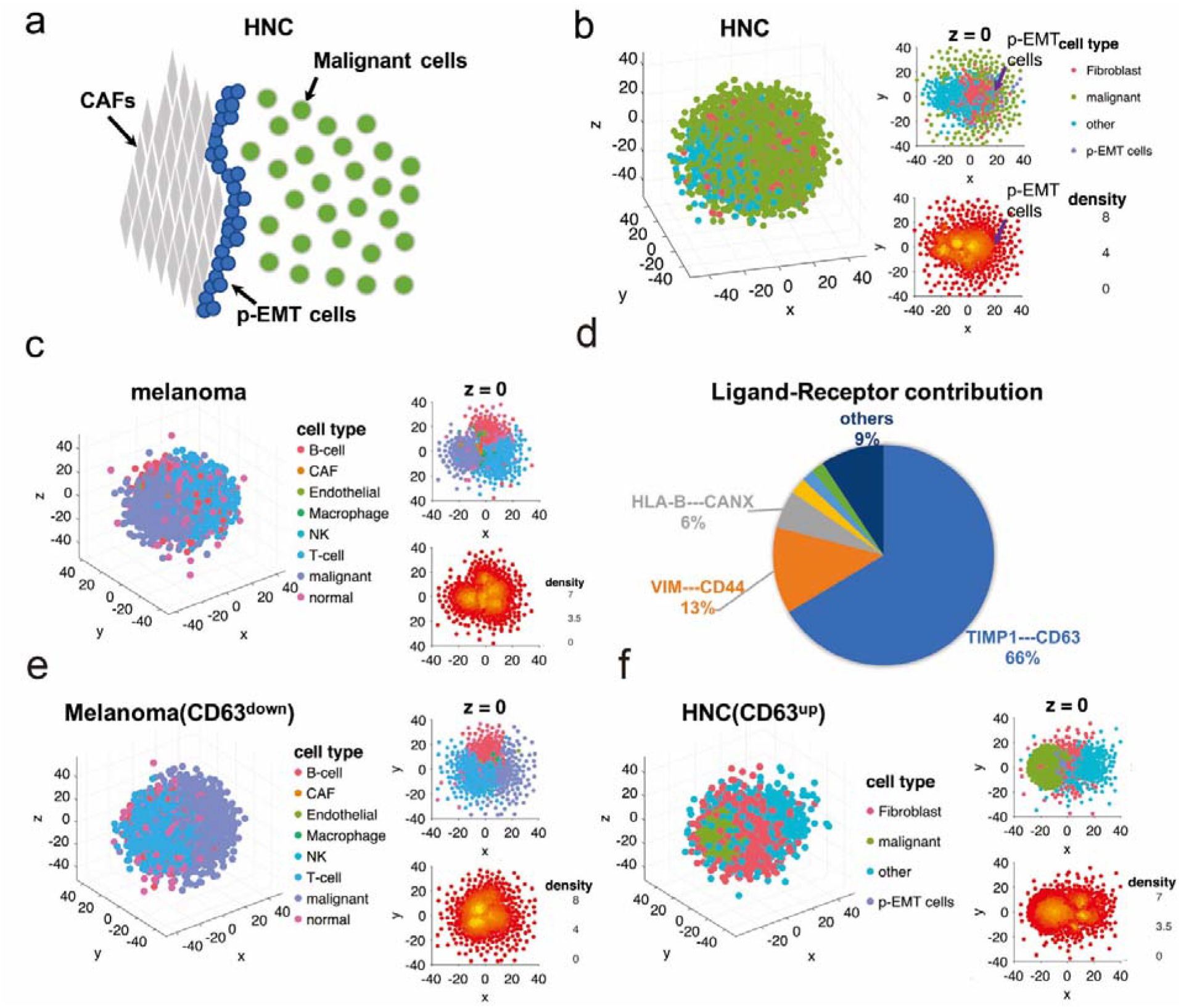
CSOmap reveals CD63-TIMP1 as a critical ligand-receptor pair in determining the spatial characteristics of HNC and melanoma malignant cells. (a) Cartoon illustration of spatial characteristics observed in IHC images of HNC patients^22^. (b) Global (left) and cross-section (right) views of reconstructed spatial organization of HNC cells. (c) Global (left) and cross-section (right) views of reconstructed spatial organization of melanoma cells. (d) Contributions of ligand-receptor pairs to the interactions among melanoma malignant cells. Spatial characteristics of melanoma (e) and HNC (f) after altering the expression levels of CD63.X

Further biological insights were also provided by the spatial reconstruction of CSOmap based on the melanoma and HNC scRNA-seq data. First, the malignant cells of both melanoma and HNC did not show significant interactions with T cells according to our CSOmap analyses, which may partially explain the immune evasion of these tumors. By contrast, while the CAFs, endothelial cells and macrophages account for much smaller fraction of the datasets, they showed statistically significant interactions with almost all the other cell types. These results recapitulate the critical roles of these cells in the spatial organization of tumors and in the regulation of tumor-infiltrating lymphocytes^25^. Second, comparison of the spatial organizations between melanoma samples that are treatment-naïve^23^ and on immunotherapy^26^ highlights the tumor and T cell compartments observed by IHC. Upon treatment, increased tumor-T interactions were observed by IHC (also revealed by CSOmap), indicating the potential effects of immunotherapy. Differential gene expression analysis based on the CSOmap prediction indicated that malignant cells not interacting with T cells show lower levels of class I major histocompatibility complex (MHC) molecules and JUN but higher level of CDK6. These results recapitulate the cancer cell program contributing to resistance of immune checkpoint blockade in melanoma identified recently^26^, suggesting the effectiveness of CSOmap in generating valid biological insights.

CSOmap additionally allows *in silico* cellular perturbation. We applied CSOmap to three scRNA-seq datasets based on T cells from the peripheral blood, tumors and tumor-adjacent normal tissues of patients with hepatocellular carcinoma (HCC)^27^, non-small cell lung cancer (NSCLC)^28^, or colorectal cancer (CRC)^29^. Since tumor- and normal-tissues and blood have distinct morphologies, and that tertiary lymphoid structures (TLSs) are frequently found in tumors^30^, we hypothesize that T cells infiltrating into different tissues may also demonstrate distinct spatial organization characteristics. CSOmap analyses on all the three datasets suggest that T cells from tumors tend to have significantly more interactions with themselves while T cells from the peripheral blood tend to disperse from each other (Supplementary Fig. 6a), confirming our hypothesis. Cellular density analysis clearly indicated the existence of tightly-linked structures (Supplementary Fig. 6b), with tumor-derived T cells forming the major part of these structures (Supplementary Fig. 6c). It has been reported that Tregs tend to trap tumor-infiltrating CD8+ T cells into TLSs or draining lymph nodes^31^. Consistently, our analysis indicated that Tregs and tumor-infiltrating Texs are the major parts of these compact structures (Supplementary Fig. 6d), which we speculate to correspond to TLSs in or near tumors. Interestingly, although blood-derived T cells did not show significant interacting potential to each other compared to tumor-derived T cells, those compact structures composed of blood-derived T cells were also observed in the HCC and NSCLC datasets, supporting the colony-forming capacity of T lymphocytes from peripheral blood as reported previously^32^.

Among T cells, Tregs and Texs exhibited significant interaction, with the ligand-receptor pair CCL4-CCR8 driving such interaction (Supplementary Fig. 6e). While CCL4 was highly expressed in most activated CD8+ T cells, its expression level in Texs was two-fold of that in other cells. CCR8 was specifically and highly expressed in tumor Tregs. According to the spatial organization reconstructed by CSOmap, Texs could be further divided into two subgroups: Texs interacting and not interacting with Tregs. Consistent with a previous report^33^, MKi67 levels were depleted in Texs interacting with Tregs (Supplementary Fig. 6f), suggesting inhibited proliferation by Tregs. Since Texs are characterized by high expression of T cell exhaustion markers including PDCD1, CTLA4, HAVCR2, TIGIT and LAG3, we examined the expression levels of their ligands in Tregs. *CD274*+ (or *PDL1*, gene for the PDCD1 ligand) and *CD80*+ (gene for CTLA-1 ligand) Tregs were significantly enriched in Tregs interacting with Texs while CD86+ Tregs were enriched in Tregs not interacting with Texs in CRC (Supplementary Fig. 6g). These results suggest that Tregs might suppress CD8+ T cells via PD-1 and CTLA-4-mediated co-inhibitory axes. Similar trends were also found in HCC and NSCLC despite of varying significance. In addition to Texs, Tregs also showed significant interactions with a set of CD8+ effector memory T cells (Tems). In CRC, T cell receptor (TCR)-based tracking suggests frequent state transitions between Tems and Texs in tumor^29^. The ratio of Tems to Texs in tumor has also been associated to better survival in lung adenocarcinoma patients^28^ and better response to immunotherapies in melanoma recently^34^. The notable interaction between Tregs and Tems in tumor may suggest a role of Tregs in the early stage of immune evasion of tumors. Similar to Texs, Tems interacting with Tregs showed higher levels of IFNG and TNF than those not interacting, and the expressions of IFNG and TNF showed significant correlations with CCL4 in Texs/Tems interacting with Tregs rather than those not interacting, suggesting that functional CD8+ T cells were prone to be targeted by Tregs due to high level of CCL4 secretion. IHC analysis of HCC and CRC samples confirmed the interactions of Tregs with Texs and Tems based on the colocalization of Tregs and CD8+ T cells in tumor (Supplementary Fig. 5, 6h and 7).

*In silico* Treg depletion by CSOmap revealed that a subset of blood-enriched recently activated effector memory T cells (Temras) demonstrate significantly increased interactions with Texs via the CXCR3-CCL5 axis (Supplementary Fig. 8), which is different from the CCR8-CCL4 axis mediating the Treg-Tex interactions. It has been recently reported in murine and human melanoma that, compared with CCR2 and CCR5, CXCR3 is necessary for the successful trafficking of tumoricidal T cells across tumor vascular checkpoints^35^, consistent with our finding that CXCR3 might mediate the migration of blood T cells into tumor via the CCL5 gradient. Treg depletion also increased the interactions of a CD4+CXCR6+ tissue-resident helper T cells (Ths) with Texs via the CXCR3-CCL5 axis (Supplementary Fig. 8), supporting the role of T cell competition in immune regulation revealed previously^36^.

CSOmap also enables computational simulation of adoptive cell transfer, which has proven to be an effective immunotherapy for cancer treatment^37^. It is currently difficult to experimentally evaluate the phenotypic outcome of adoptively transferred T cells. We used the HNC and melanoma datasets as foundations to simulate their tumor microenvironments and used blood- and tumor-derived T cells for *in silico* adoptive transfer. We simulated a gradient of TCR-pMHC affinity between the adoptively transferred T cells and the malignant cells and quantified the numbers of tumor-T interactions, tumor-infiltrating T cells and targeted malignant cells. Interestingly, while the numbers of interactions between T and malignant cells increased in a linear function of the TCR-pMHC affinity, those infiltrating T cells and targeted malignant cells increased in a logarithmic function (Supplementary Fig. 9). Visually, an interface formed between T cells and malignant cells (Supplementary Fig. 9). This phenomenon observed *in silico* might recapitulate and explain the morphological patterns frequently observed in tumor microenvironment by IHC and multiplexed ion beam imaging^38^. While the TCR-pMHC affinity was the dominant determinant of T-malignant cell interactions, the phenotypes of T cells and malignant cells also contributed significantly (Supplementary Fig. 9 and 10). In particular, tumor-derived T cells showed significantly higher efficiency in tumor infiltration than blood-derived T cells (Supplementary Fig. 9) while malignant cells of melanoma were more prone to be targeted than those of HNC (Supplementary Fig. 10) according to ANOVA analysis with repeated measures. These results recapitulated, from a computational perspective, the variations of T cell transfer-based therapies across cancers and the predictive values of immunophenotypic characterization of infused T cell product in engraftment and responses observed in various clinical trials^39^.

In summary, CSOmap provides a computational tool to *de novo* reconstruct cellular spatial organization from scRNA-seq data. The underlying assumption is that cells can compete and self-assemble into specific spatial patterns via ligand-receptor interactions. Despite of the incomplete nature of known ligand-receptor interactions and their affinity parameters (particularly for TCR-pMHCs), the dropout issue of scRNA-seq, biases or errors of estimating the protein-level abundance of ligands and receptors by transcriptomic data, the distinction of capacity to interact from realistic interactions, and the unavailability of other physical, chemical, and nutrient factors involving cell organizations, evaluations on multiple scRNA-seq datasets spanning human and mouse physiological and diseased conditions demonstrated that CSOmap can recapitulate critical spatial characteristics qualitatively and quantitatively. Due to the convenience of *in silico* manipulation, CSOmap can also generate profound insights into the molecular mechanisms underlying spatial formation via *in silico* simulation of gene overexpression, knock-down, cell adoptive transfer and depletion. CSOmap will be applicable to interrogating cellular organizations from scRNA-seq data for various tissues in diverse systems, and it can be greatly enhanced when more complete knowledge of ligand-receptor interactions and other critical factors are available.

## ACKNOWLEDGMENTS

We thank Boxi Kang for his IT assistance. We thank the Computing Platform of the CLS (Peking University) for providing computing resource. This project was supported by Beijing Advanced Innovation Centre for Genomics at Peking University, Key Technologies R&D Program (2016YFC0900100), and National Natural Science Foundation of China (31991171, 31530036 and 91742203).

## AUTHOR CONTRIBUTIONS

X.R. and Z.Z. conceived this study. X.R. designed the mathematical model and algorithm. G.Z. developed the software. X.R. and G.Z. performed the analysis. Q. Z., L.Z. and Y. S. conducted the validation experiment. X.R., G.Z. and Z.Z. wrote the manuscript with all the authors’ inputs.

## ONLINE METHODS

### Overview of CSOmap

CSOmap reconstructs cellular spatial organization of individual cells from single-cell RNA sequencing data based on three principles: 1) Cellular spatial organization is determined by ligand-receptor mediated cellular self-assembly, with cells having high affinity close to each other spatially; 2) The affinity of cells can be defined by the abundance of ligands and receptors and their interacting potentials; 3) Cells compete with each other to form spatial structures. Hence, the core algorithm of CSOmap to reconstruct spatial organization of single cells from RNA sequencing data includes two steps: 1) estimating the cellular affinity matrix based on the gene expression profiles of individual cells and known ligand-receptor interactions; 2) embedding the inherently high-dimensional cellular affinity matrix into three-dimensional pseudo-space resembling the realistic biological tissues during which cell competitions are sufficiently considered. CSOmap also includes additional algorithms for analyzing the resultant three-dimensional coordinates of single cells, including estimating the density of each cell, identifying spatially-defined cell clusters/structures, evaluating the number of connections and statistical significance between two cell clusters defined by expression profiles or other characteristics, calculating the contributions of each ligand-receptor pair to the interaction potential of two cell clusters, and *in silico* molecular and cellular interference. The details of the core and additional algorithms of CSOmap are depicted as follows separately.

### Estimating cell-cell affinity by ligand-receptor interactions

The estimation of cell-cell affinity is critical to the performance of CSOmap. To define a valid function of cell-cell affinity, we assume that the affinity of two cells equals to the affinity summation of all the protein complexes forming by the proteins from the surfaces or extracellular matrices of two cells. We applied a series of approximations to facilitate computation at the genome scale. Details are stated as follows.

In biological reality, the number of components of a protein complex varies from two to tens. For computational convenience, we converted all the interactions of more than two components to binary interactions without regarding the complicated nonlinear effects. Given a binary interaction, *i.e.*, one ligand A and one receptor B, according the law of mass action in chemistry, the concentration of the complex AB can be calculated according to the following formula:

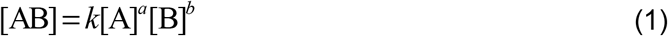

where [AB] is the concentration of the complex AB, [A] is the concentration of the ligand A, [B] is the concentration of the receptor B, *k* is the reaction constant, and *a* and *b* are the stoichiometric coefficients of A and B, respectively. The parameters *k*, *a* and *b* vary according to the chemical nature of A and B. For similarity, we approximate Formula (1) by the following formula to handle thousands of pairs of ligand and receptor:

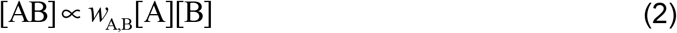

where *W*_A,B_ is introduced to summarize the total effects of the parameters *k, a* and *b*. Upon this approximation, the cell-cell affinity is defined by the following formula:

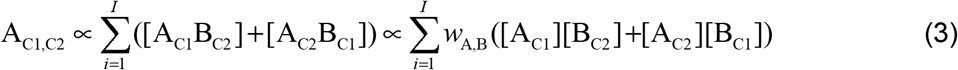

where A_C1,C2_ denotes the affinity of Cell C1 and Cell C2, [A_*c*_] or [B_*c*_] denotes the concentration of the A or B molecule on Cell *c*, *i*is the index of ligand-receptor pairs, and there are a total of *I* pairs. Because the ligand and receptor can be simultaneously expressed by both of the cells, a symmetric term of the concentrations of the complex is added. Furthermore, we use the mRNA abundance of the ligand and receptor to approximate their protein concentrations, and thus Formula (3) can be updated as follows:

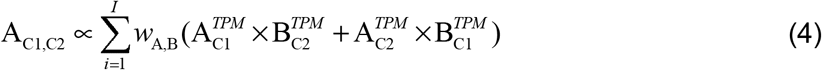

where 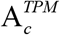 or 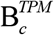 is the mRNA level of the ligand A or receptor B in Cell *c* estimated by the Transcripts Per Million (TPM) measure. Since there are thousands of ligand-receptor interactions, for which most of the parameters of their interacting dynamics (summarized by *W*_A,B_) are not available, we set *W*_A,B_ = 1 in the current version of CSOmap while providing *W*_A,B_ as a parameter of the software for incorporating users’ knowledge of the chemical natures of the ligand-receptor interactions. According to (4), it is thus established of the computational method for estimation of cell-cell affinity based on single-cell RNA sequencing data and ligand-receptor interactions. In practice, we used the human ligand-receptor interaction database FANTOM5 with incorporation of immune-relevant chemokines, cytokines, co-stimulators, co-inhibitors and their receptors for estimating the cell-cell affinity matrix^14, 40^ (Supplmentary Table 1). Some of these ligands such as chemokines and cytokines are not membrane-located but secreted proteins. We included these ligands into the estimation of cell-cell affinity similar to those membrane proteins because they often form gradients to affect the migration of other cells, particularly for chemokines. Interactions involved in B2M were manually filtered because of its housekeeping nature. Ligand-receptor interactions in the CellphoneDB^18^ database were also included to show the robustness of CSOmap predictions (Supplementary Table 2). Because of the potential noises introduced in the estimation of cell-cell affinity due to noises in gene expression levels and various approximations, we further discretize the cell-by-cell affinity matrix by retaining the top *k* highest-affinity neighbors for each cell to reduce noise (k=50 by default).

### Embedding the high-dimensional cell-cell affinity matrix into three-dimensional space

When the discretized cell-by-cell affinity matrix is obtained, this inherently high-dimensional matrix is further embedded into a three-dimensional space. The principle behind this operation is that the realistic biological space is of only three dimensions and that positive affinity values in the affinity matrix only indicate the interacting potentials rather than realistically occurred facts. Cells of interacting potentials with common targets need to compete the space to change potentials to reality. For considering this factor, we introduce three constraints to build the computational model. First, a minimum distance between cells is pre-defined because all cells have positive sizes and cannot be ultimately squeezed. Second, the total available space is also pre-defined by a parameter named as space radius to simulate the limited realistic space. Finally and the most importantly, Student’s t-distribution is introduced to resolve the crowding issues of cell-cell interactions, motivated by the visualization algorithm t-SNE, which allows cells to compete with others to form the spatial organization. Taken all these considerations together, we propose the following computational model as the core of CSOmap:

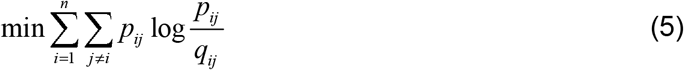

subject to:

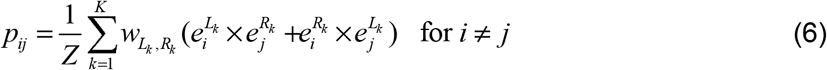

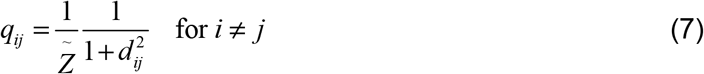

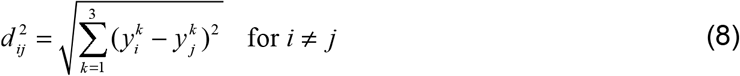

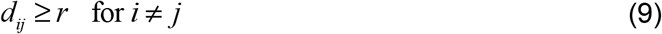

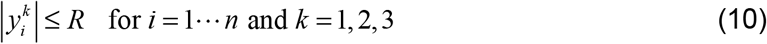

where 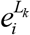 or 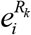 is the TPM values of the *k*-th ligand or receptor in the *i*-th cell, *w_L_k_, R_k__* is the weight summarizing the chemical nature of the *k*-th pair of ligand-receptor, *p_ij_* is the cell-cell affinity between *i* and *j* estimated by the aforementioned method, 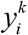 is the *k*-th coordinate of the *i*-th cell, *d_ij_* is the Euclidean distance between the *i*-th and *j*-th cells in the embedded space, and *q_ij_* is the probability of the *j*-th cell locating in the neighborhood of the *i*-th cell. Constraints (6)–(8) give out the definitions of *p_ij_*, *q_ij_*, and *d_ij_* while constraints (9)(10) impose the spatial limitations. Kullback-Leibler divergence is used to define the loss function (5). This optimization model is highly similar to the model used to implementing non-linear dimensional reduction in the frequently used visualization algorithm t-SNE except that constraints (9)(10) are imposed to consider the spatial limitations. Similarly, a gradient-descent algorithm is used to solve this optimization problem with random initialization and then updating the solution with the guidance of the gradient. When the maximum number of iterations was reached, the resultant three-dimensional coordinates were reported for subsequent analyses. In principle, large *r* and small *R* will reduce the volume of cell space and thus provide repulsive forces while the cell-cell affinity provides attracting forces. The repulsive and attracting forces together guide the self-assembly of cells and finally determine the cellular spatial organizations. Particularly, large *r* and small *R* will introduce fluctuations into the cellular spatial organizations. To obtain a stable organization, we set *r* =0.1 and *R* =50 in practice. The initializing solution is randomly assigned in a 50×50×50 cube, and the maximum number of iterations is set to 1000. When the three-dimensional coordinates are obtained, a rotation of the coordination system is made by principal component analysis to guarantee the X and Y axes capturing the most spatial variations. By default, we set the number of dimensions as 3 because biological tissues/organs are in 3D space, but the users can tune this parameter to 2 or 1 to examine specific tissue models.

### Density analysis and clustering spatially compact cell clusters

Given the three-dimensional coordinates of all cells, a straightforward analysis is to check what spatial structures are formed, which can be examined visually and quantitatively. CSOmap implements a series of visualization method to facilitate the recognition of spatially organized structures, including global three-dimensional views with various rotation angels, cross-section views with various slicing methods, and even dynamic views to show how cells self-assemble into organizations via ligand-receptor medicated interactions. Categorical or numerical features can be used to color cells to highlight the patterns. Quantitatively, the compactness of the neighborhood of a cell, named as density, can be calculated by counting the number of cells within a predefined radius. By default, the radius is set to the median distance of each cell to its third nearest neighbor because the number of neighbors of a cell cannot be too large due to limited space. When the density of individual cells are defined, clustering based on fast searching and finding density peaks^16^ can then be applied to identify spatially compact cell clusters and dissociative cells. Sensitivity analysis suggested that the identification of compact structures is robust to the selection of the radius.

### Evaluating the statistical significance of cellular interactions between cell clusters

The resultant three-dimensional coordinates of CSOmap also allow examining whether two cell clusters tend to interact with each other significantly and thus locate close to each other. Given a threshold defining the neighborhood radius of a cell, *e.g*, the median of the third nearest distance, a pair of cells can be assumed to “directly” interact with each other if their distance is less than the cutoff. Therefore, the total number of cell-cell interactions between two clusters can be counted. The statistical significance of the observed number of cell-cell interactions can further be evaluated by random permutation of the cluster labels of individual cells. With 1000 random permutations, the distribution of the randomly expected cell-cell interaction numbers of the given two cell clusters can be constructed, and thus right-tailed and left-tailed tests can be conducted respectively to calculate the p-values for the hypotheses that the observed interaction number was larger than that randomly expected (enrichment) and that the observed number was smaller than that randomly expected (depletion). When p-values for enrichment between all clusters are obtained, the Benjamini-Hochberg procedure is used to estimate the q-values. If the enrichment (depletion) q-value for a given pair of cell clusters is less than 0.05, these two cell clusters are claimed to significantly interact with (disperse away from) each other. Otherwise, if the enrichment and depletion q-values are both larger than 0.05, the cell-cell interactions are assigned to the “other” type.

### Determining the dominant ligand-receptor pairs underlying cell-cell interactions

Given a pair of cells, the contribution of a specific ligand-receptor interaction to the cell-cell interacting affinity can be calculated by the following formula:

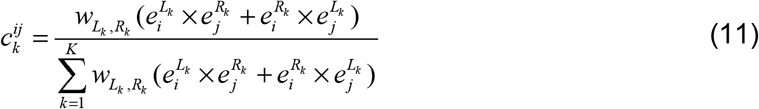

where 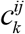 denotes the contribution of the *k*-th ligand-receptor interaction to the cell-cell affinity of the *i*-th and *j*-th cells. Therefore, given a pair of cell clusters, the contribution of a specific ligand-receptor pair to the interactions of the two clusters can be calculated by the following formula:

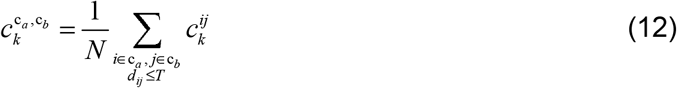

where 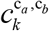 denotes the contribution of the *k*-th ligand-receptor interaction to the interactions of two clusters c_*a*_ and c_*b*_, and *N* is the total number of cell pairs between c_*a*_ and c_*b*_ that conform to the definition of “direct” cell-cell interaction, *i.e.,* the distance between two cells *i* and *j* should be less than the predefined threshold *T* (*d_ij_* ≤ *T*). When the contributions of all the ligand-receptor pairs are calculated, the ligand-receptor pairs with the highest scores are assumed to be the dominant molecular contributors underlying the cell clusters.

### Evaluating the spatial effects of individual genes or cell clusters by *in silico* interference

CSOmap also provides functions to analyze the effects of *in silico* interfering genes or cell clusters on the cellular organization. In reality, cell-cell interactions form a highly nonlinear system, and thus it is hard to predict the spatial effects of gene alterations or cell interference. By simulating the cellular spatial organization via ligand-receptor mediated self-assembly, CSOmap provides an easy way to interrogate the nonlinear effects of ligand/receptor or cellular changes on the cellular organizations, and thus can provide important insights into the true biological mechanisms that are too expensive or even impossible to obtain by experimental methods. Currently, the *in silico* interference types of CSOmap include *in silico* gene knockdown, gene overexpression, adoptive cell transfer, and cell depletion. When cellular spatial organization with *in silico* interference is obtained, it can then be compared to the original organization to identify the significant differences. Although the current *in silico* interference can only examine the effects of ligand and receptors, it can be further enhanced by incorporating gene-gene interactions in the future to introduce dynamics for the expression levels of ligands and receptors. Since the output coordinates of CSOmap are in virtual spaces, it is now not possible to directly compare the changes of cellular spatial organizations at the coordinate level. All the comparisons in the current manuscript were conducted after abstracting the coordinate results into cell-cell interacting graphs. For the HNC dataset, *in silico* CD63 overexpression was conducted through changing all the original CD63 expression values to TPM 5000. For the melanoma dataset, in silico CD63 knockdown was implemented by resetting the original expression values to zero. For the CRC T cell dataset, Treg depletion was implemented by removing all the cells belonging to the CD4-CTLA4 cluster.

### Performance evaluations of CSOmap

The mouse liver lobule and paired-cell sequencing datasets were downloaded from the Gene Expression Omnibus database (https://www.ncbi.nlm.nih.gov/geo/) with accession numbers GSE84498 and GSE108561, respectively. The human and mouse pancreas datasets were downloaded with the access number GSE84133. The 10X Genomics-based human placenta dataset was downloaded from http://data.teichlab.org (maternal–fetal interface). The 10X Genomics-based Tabula Muris datasets were downloaded with the access number GSE109774, with neuron and immune cells removed to investigate the interactions within and between organs.

The HNC and melanoma (TN) datasets were downloaded with access numbers GSE103322 and GSE72056, respectively. The melanoma (ICR) dataset was downloaded through the Single Cell Portal (https://portals.broadinstitute.org/single_cell/study/melanoma-immunotherapy-resistance). The T cell datasets of HCC, NSCLC and CRC were downloaded from the Gene Expression Omnibus with accession numbers GSE98638, GSE99254 and GSE108989. All the expression values of the original datasets were converted to TPM for CSOmap analysis. The reconstructed spatial organizations of HNC and melanoma datasets were compared to the IHC images published in their original papers by examining the spatial characteristics of various cell groups. The reconstructed spatial organizations of the three T cell datasets were validated by comparing their biological corollaries with published literature. Because cellular spatial organization is the result of many factors including cell identity, cellular environment, cell developmental history, and many physical and chemical ingredients, it is currently impossible to directly compare the precise structures of the reconstructed organizations with experimentally obtained images. Comparison of the spatial characteristics and biological corollaries is acceptable. The IHC and scRNA-seq experiments of the HCC sample was conducted in house, with the scRNA-seq operations following the protocol of SMART-seq2 and the IHC experiment and image analysis following the protocols provided by Nghiem et al. 2016^41^ and the PerkinElmer Vectra automated multispectral microscope. The scRNA-seq data were deposited into EGA with access id EGAS00001003449. The antibodies used in the IHC staining were from Abcam: PD1: [EPR4877(2)] (ab137132); CD8: [144B] (ab17147); CD68:[EPR20545](ab213363); FOXP3: [236A/E7](ab20034); CLEC9A: [8F9](ab104910).

### T cell adoptive transfer analysis

In the *in silico* T cell adoptive transfer experiments, the HNC and melanoma (TN) datasets were used to simulate the corresponding tumor microenvironments. For fair comparison, the adoptively transferred T cells were sampled from the HCC T cell dataset. For each *in silico* adoptive transfer experiment, the number of adoptively transferred T cells were set to the same as the number of T cells of the original datasets. To simulate the TCR-pMHC interactions between adoptively transferred T cells and the malignant cells, a pair of pseudo-ligand and receptor was added into the ligand-receptor network. The pseudo ligand (pMHC) was set to be only expressed in malignant cells with TPM 5000 and the pseudo receptor (TCR) was set to be only expressed in adoptively transferred T cells with varying expression levels. The TCR-pMHC affinity was represented by the varying levels of the pseudo-receptor. ANOVA with repeated measures (implemented by the ranova function of Matlab R2016b) was used to evaluate the effects of TCR-pMHC affinity, origins of T cells, and cancer types.

### Data availability

The mouse liver lobule and paired-cell sequencing datasets were downloaded from the Gene Expression Omnibus database (https://www.ncbi.nlm.nih.gov/geo/) with accession numbers GSE84498 and GSE108561, respectively. The human and mouse pancreas datasets were downloaded with the access number GSE84133. The 10X Genomics-based human placenta dataset was downloaded from http://data.teichlab.org (maternal–fetal interface). The 10X Genomics-based Tabula Muris datasets were downloaded with the access number GSE109774, with neuron and immune cells removed to investigate the interactions within and between organs. The HNC and melanoma (TN) datasets were downloaded with accession numbers GSE103322 and GSE72056. The melanoma (ICR) dataset was downloaded through the Single Cell Portal (https://portals.broadinstitute.org/single_cell/study/melanoma-immunotherapy-resistance). The T cell datasets of HCC, NSCLC and CRC were downloaded from the Gene Expression Omnibus with accession numbers GSE98638, GSE99254 and GSE108989. The newly added HCC scRNA-seq data were deposited into EGA with access id EGAS00001003449.

### Software availability

The software implementation of our CSOmap method is available at https://doi.org/10.24433/CO.8641073.v1.

---

**Supplementary Fig. 1.**
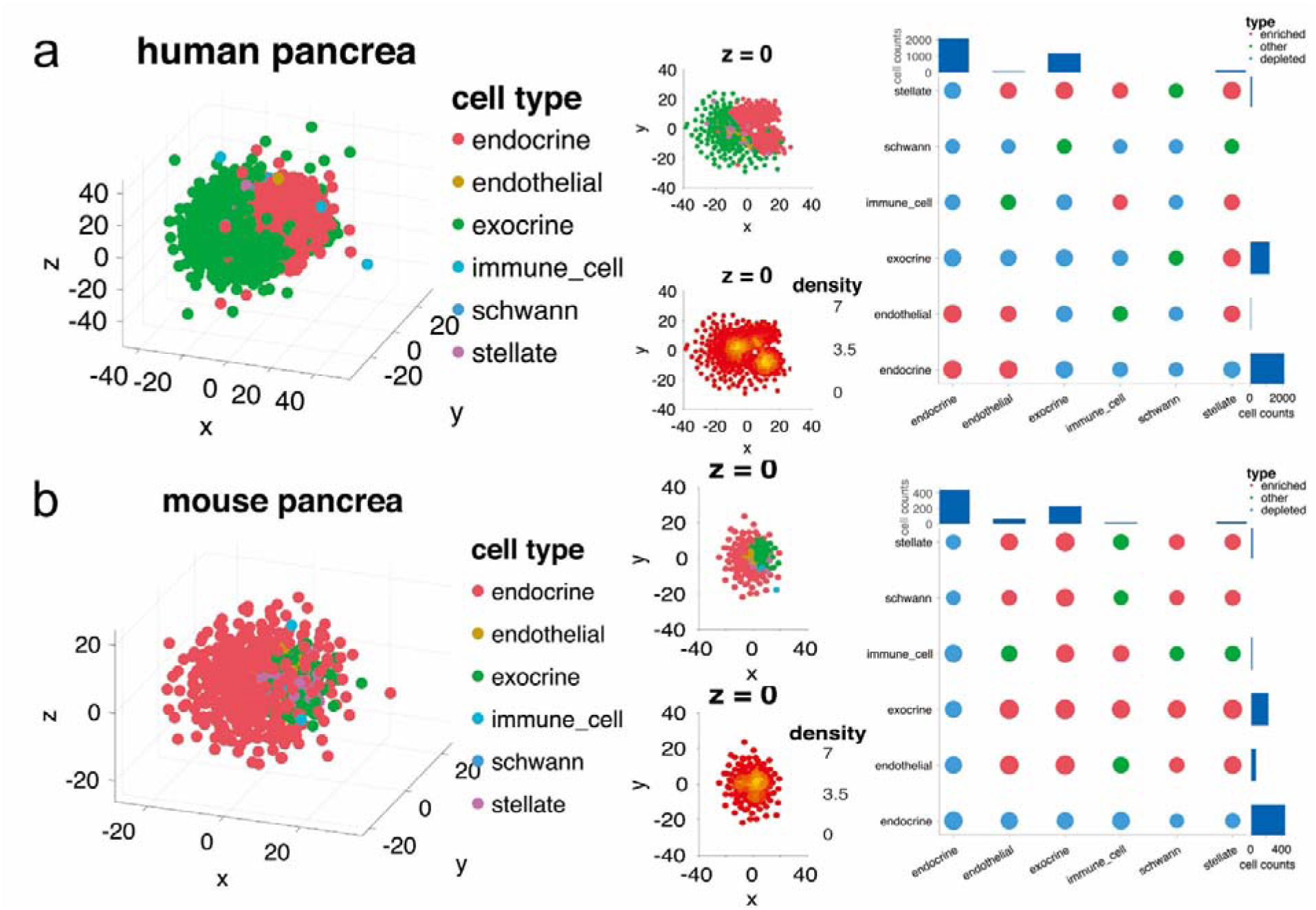
The exocrine and endocrine compartments of pancreas can be recapitulated by ligand-receptor based inference with CSOmap. a, the 3D visualization of CSOmap prediction of the human pancreatic scRNA-seq data (left), the cross-section of z=0 of the 3D visualization (middle), and the statistical significance of interactions between different cell types (right). b, the 3D visualization of CSOmap prediction of the mouse pancreatic scRNA-seq data (left), the cross-section of z=0 of the 3D visualization (middle), and the statistical significance of interactions between different cell types (right). Enriched: cells of one cell type are enriched in the neighborhood of the other cell type, p (right tail) < 0.05 and q < 0.05; depleted: cells of one cell type are depleted in the neighborhood of the other cell type, p(left tail) < 0.05 and q < 0.05. Exocrine: acinar and ductal cells; endocrine: alpha, beta, gamma, delta and epsilon cells.

**Supplementary Fig. 2.**
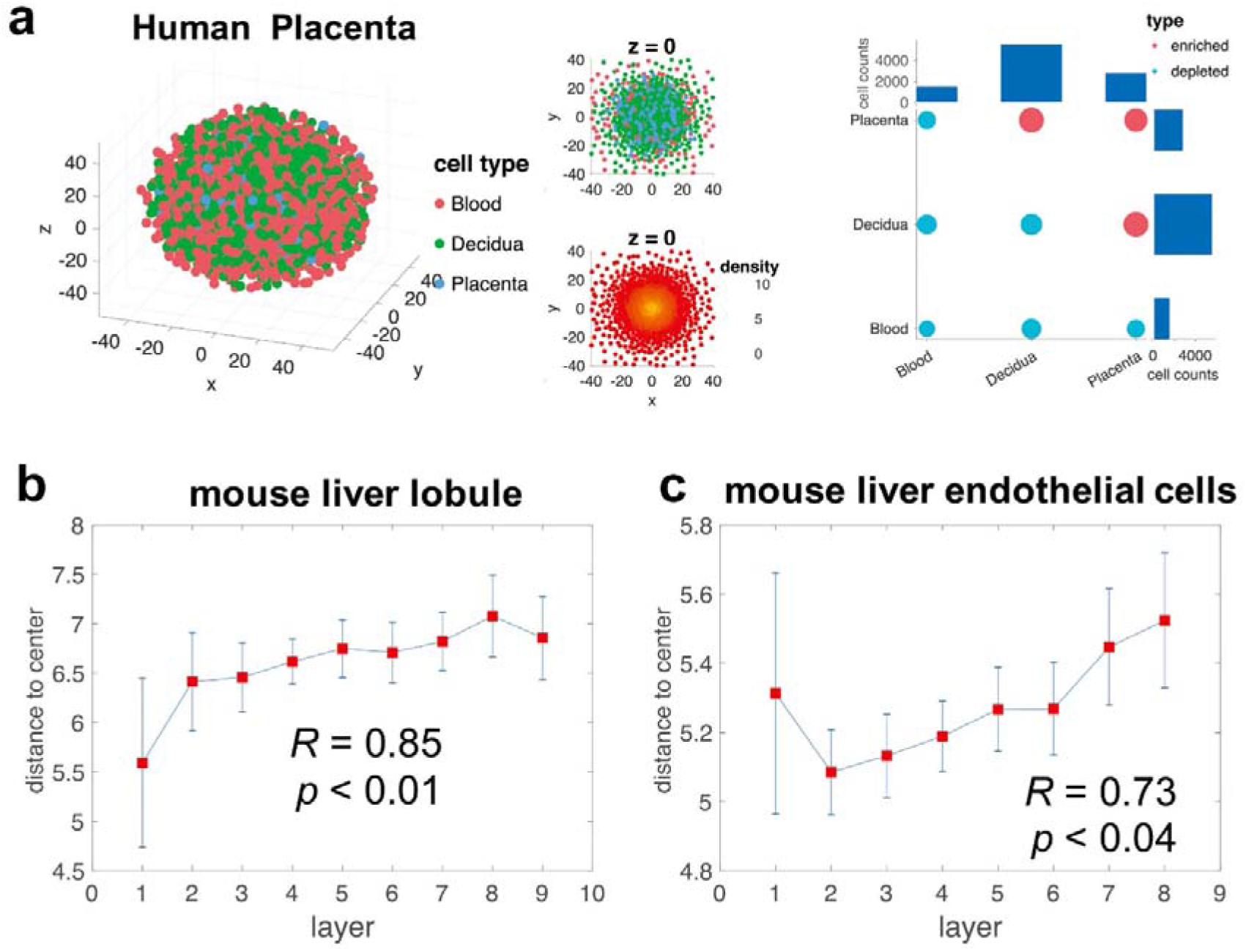
Performance of CSOmap on scRNA-seq data of human placenta and mouse liver lobules. a, the 3D visualization of CSOmap prediction based on the 10X Genomics data of the human placenta scRNA-seq data (left), the cross-section view, and the statistical significance (right). Significant interactions between fetal placenta cells and maternal decidua cells rather than blood cells can be observed, resembling the maternal-fetal interface reported in the original paper^18^. b, the consistence of CSOmap prediction with the annotated cell layer information (Pearson correlation) of a scRNA-seq data for mouse liver lobules^8^. c, the consistence of CSOmap prediction based on the paired-cell sequencing data with the annotated layer information of mouse liver endothelial cells (Pearson correlation)^19^. Distance to center: Euclidean distance. Error bars: standard error. High error bars were caused by the uncertainty of the reference labels (>93% cells having uncertainty more than 2 layers estimated by the given probabilistic distributions).

**Supplementary Fig. 3.**
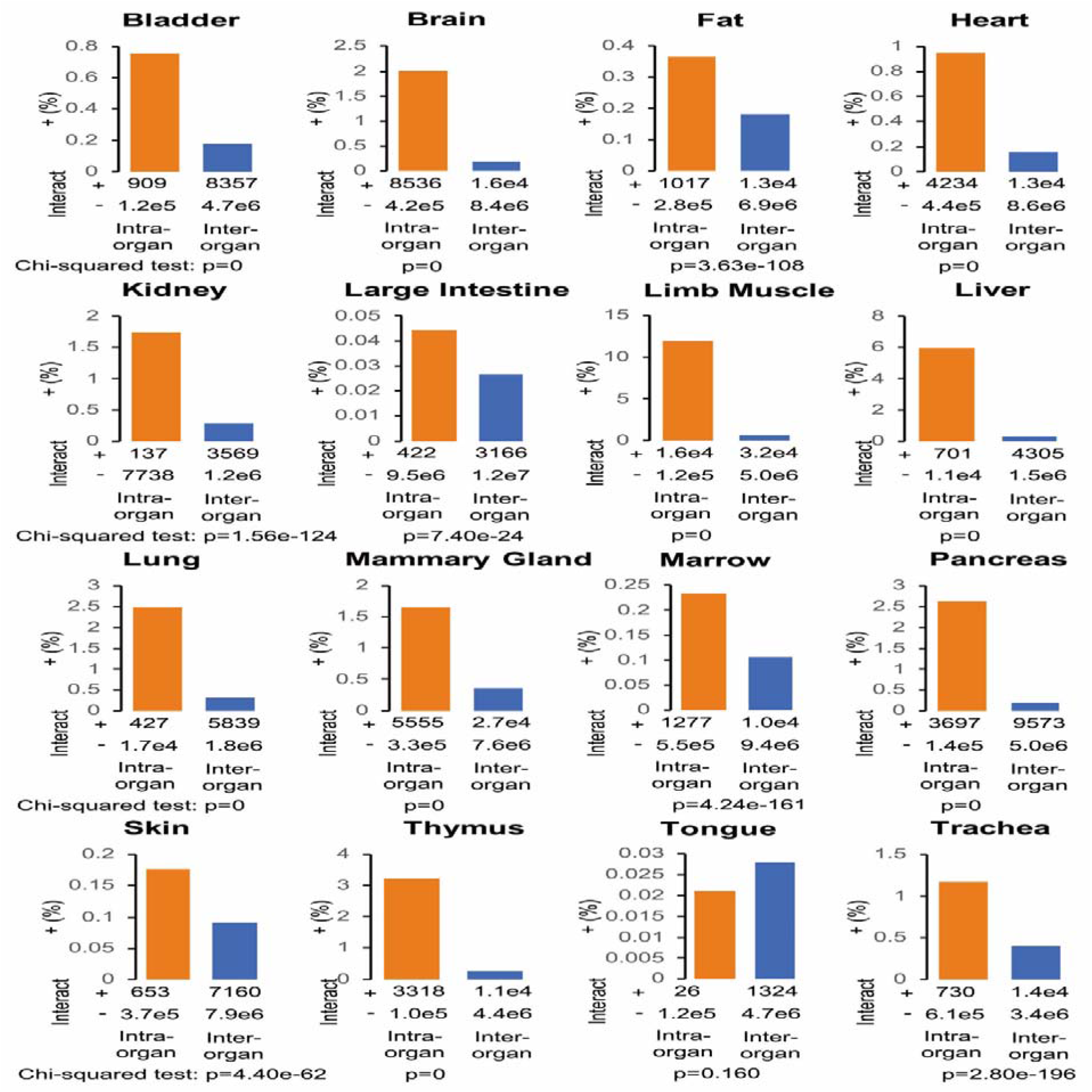
CSOmap recapitulates high intra-organ cellular interactions based on the Tabula Muris dataset. For each organ, cell pairs were categorized into intra-organ interacting pairs (Interact+Intra-organ+), intra-organ non-interacting pairs (Interact-Intra-organ+), inter-organ interacting pairs (Interact+Inter-organ+), and inter-organ non-interacting pairs (Interact-Inter-organ+), then chi-squared test was applied to examine the statistical significance of whether intra-organ cell pairs tend to have higher interacting odds than inter-organ cell pairs. The 10X Genomics data were used for CSOmap prediction. Immune and neuron cells dispersing in various organs were excluded. 16 organs were retained and 15 were predicted to have higher intra-organ cellular interactions than inter-organ interactions.

**Supplementary Fig. 4.**
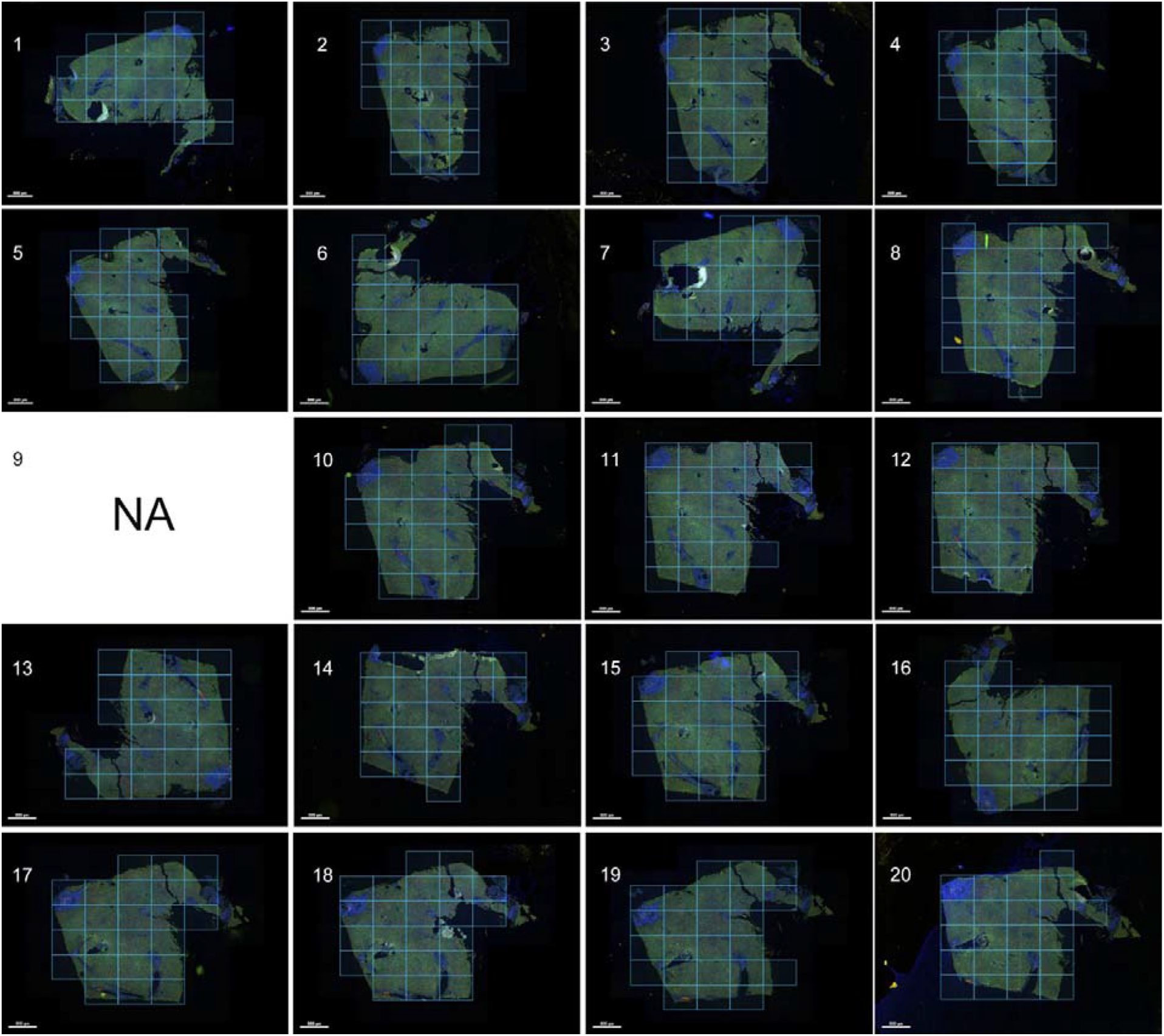
IHC images of a centimeter-scale tumor sample from hepatocellular carcinoma. 20 consecutive 1cm × 1 cm × 5 um IHC images with the 9^th^ staining failed. Treg: regulatory T cells (Foxp3+); Tex: exhausted T cell (PD-1+); CD8: CD8+PD-1^-^ T cells; cDC1: CLEC9A+ dendritic cells; M: macrophages (CD68+); O: other cells. The median distance of the 3^rd^ nearest neighbor of all cells was used as the cutoff to determine whether two cells were spatially connected or not.

**Supplementary Fig. 5.**
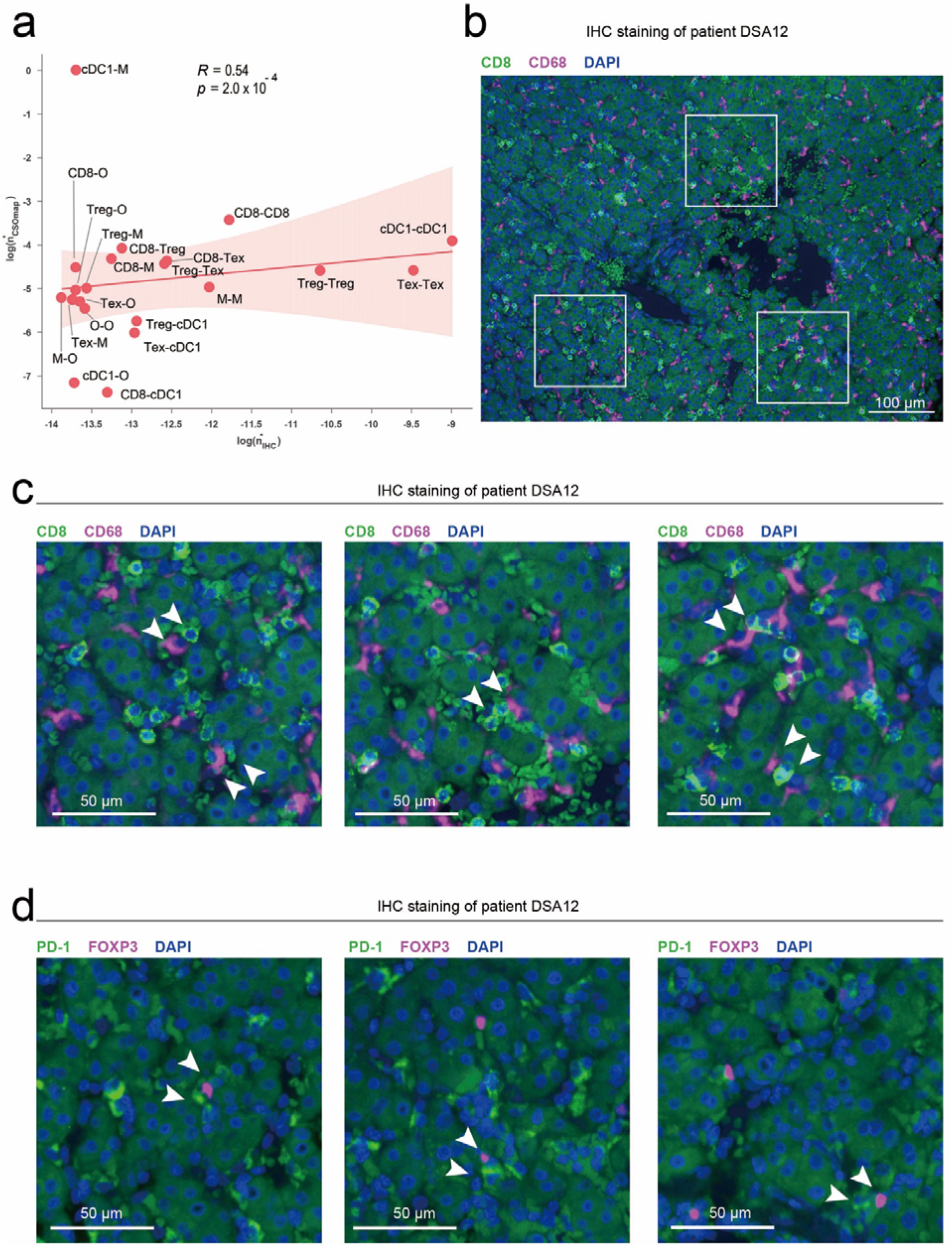
Consistence between CSOmap prediction and IHC imaging of the tumor sample from hepatocellular carcinoma. a, Spearman correlation between cell connections based on IHC images (X-axis) and the CSOmap prediction (Y-axis) after normalizing the biases introduced by uneven cell counts among different cell types. Treg: regulatory T cell (Foxp3+); Tex: exhausted T cell (PD-1+); CD8: CD8+PD-1^-^ T cells; cDC1: CLEC9A+ dendritic cells; M: macrophages (CD68+); O: other cells. b, example IHC image showing the interaction between CD8+PD-1^-^ T cells and macrophages. c, enlarged illustration of the selected windows in b (from left to right in order). d, example IHC images showing the interaction between Texs and Tregs.

**Supplementary Fig. 6.**
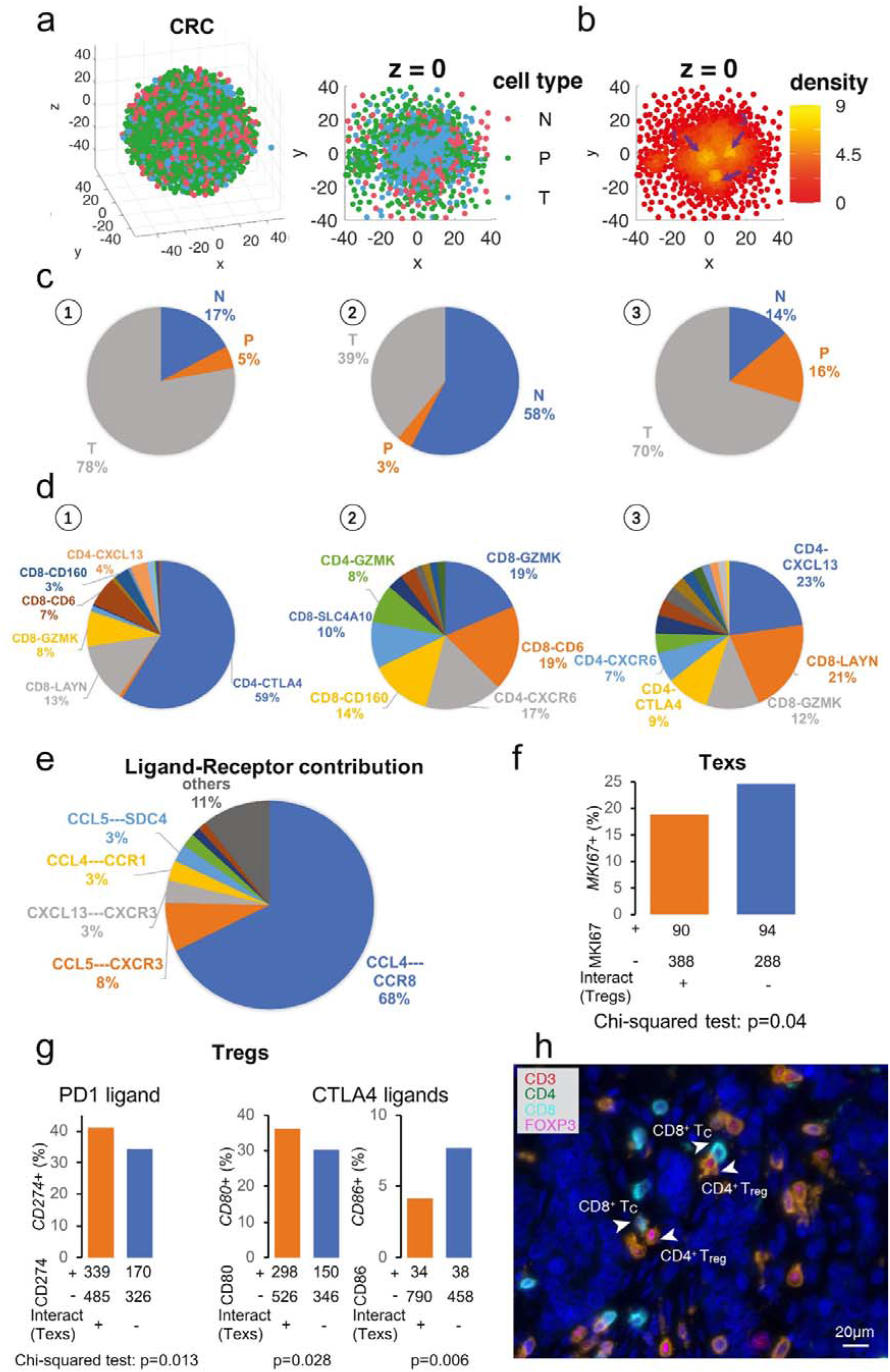
Spatial and molecular characteristics of CRC T cells. (a) Global (left) and cross-section (right) view of the reconstructed spatial organization of CRC T cells (colored by the tissue origins, N: normal tissue; P: peripheral blood; T: tumor). (b) Compactness of individual cells estimated by density, with 1, 2, and 3 indicating three observed compact structures. (c) Tissue compositions of the three observed compact structures in (b). (d) Cluster compositions of the three observed compact structures in (b), where the cluster names follow the nomenclature of the original paper with CD4-CTLA4 indicating tumor Tregs, CD8-LAYN indicating Texs, and CD8-GZMK indicating Tems. (e) Contributions of ligand-receptor pairs to the interactions between Tregs and Texs. (f) Texs interacting with Tregs showed depleted MKi67+ cells. (g) Tregs interacting with Texs showed enrichment of *CD274*+ and *CD80*+ cells and depletion of *CD86*+ cells. (h) Co-localization of Tregs and CD8+ T cells in CRC tumor samples revealed by IHC staining.

**Supplementary Fig. 7.**
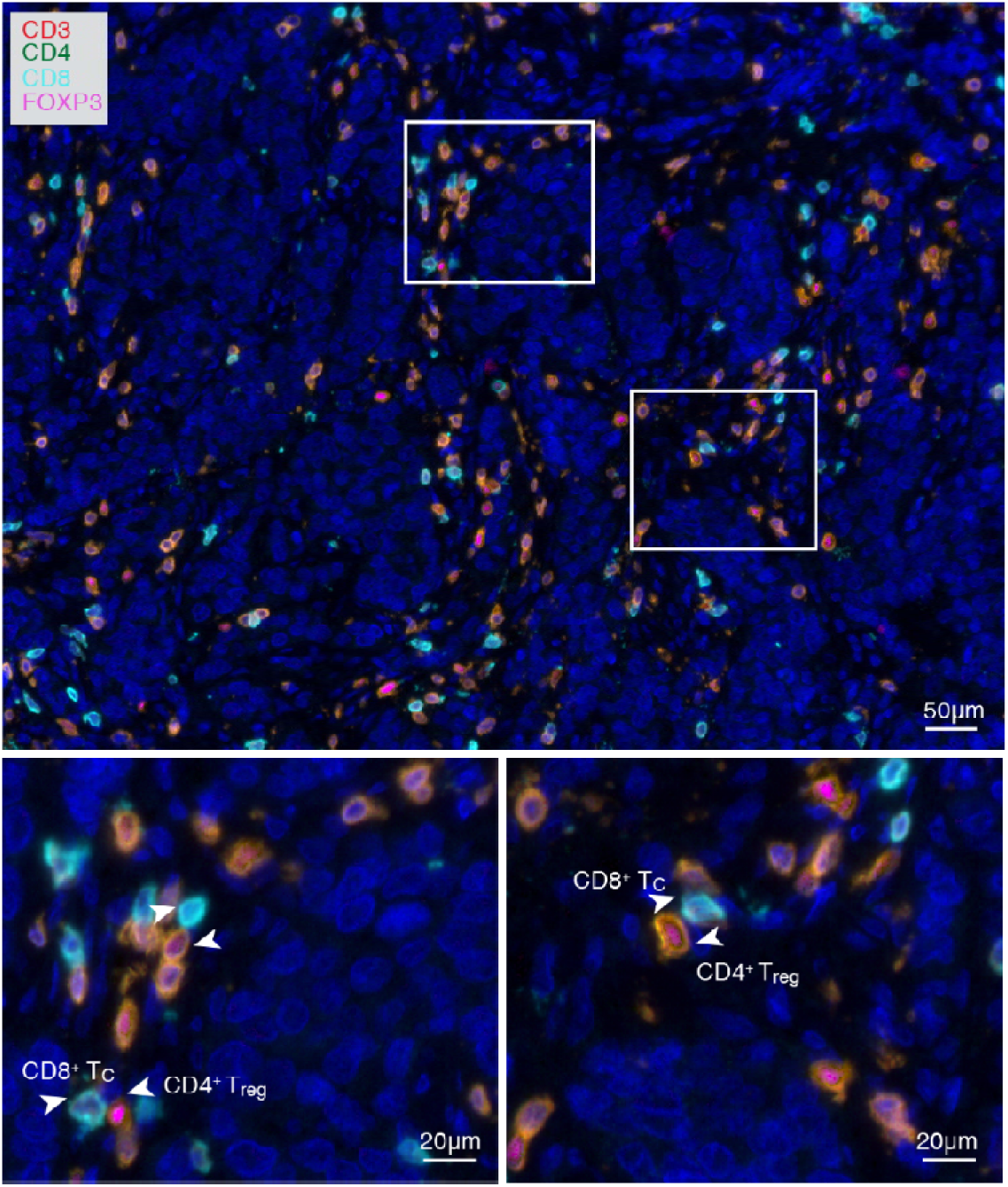
Co-localization of Tregs (CD4+FOXP3+) and CD8+ T cells revealed by IHC staining of a CRC tumor sample.

**Supplementary Fig. 8.**
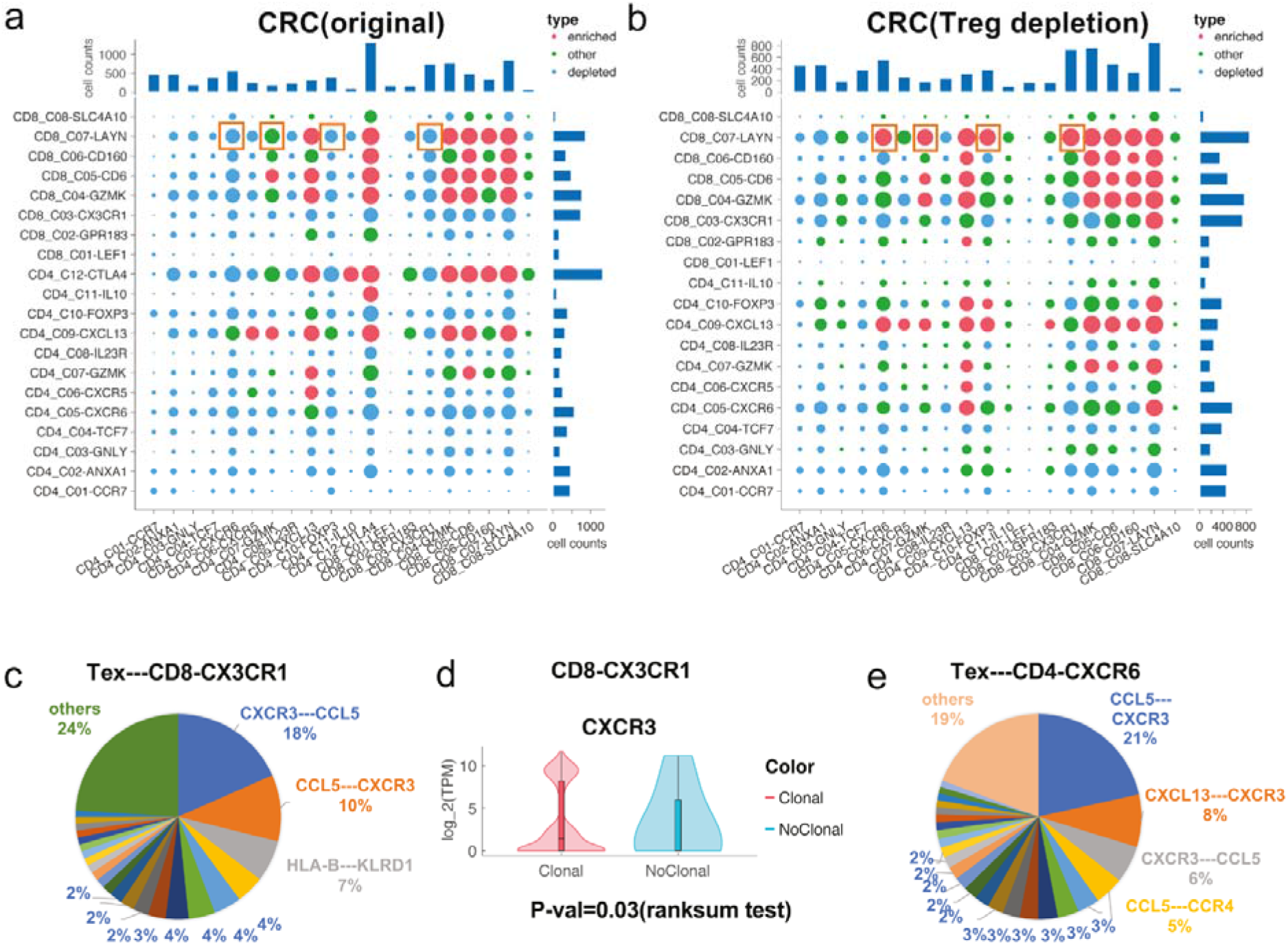
Treg depletion enhanced the interactions of Texs with blood-enriched CD8-CX3CR1 and tissue-resident CD4-CXCR6 T cells revealed by CSOmap with *in silico* interference. (a) Statistical significance of cell-cell interactions of the original dataset of CRC. (b) Statistical significance of cell-cell interactions of the CRC dataset after Treg depletion. (c) The contributions of ligand-receptor pairs to Tex—CD8-CX3CR1 interactions. (d) CXCR3 was highly expressed in clonal CD8-CX3CR1 cells. (e) The contributions of ligand-receptor pairs to Tex—CD4-CXCR6 interactions.

**Supplementary Fig. 9.**
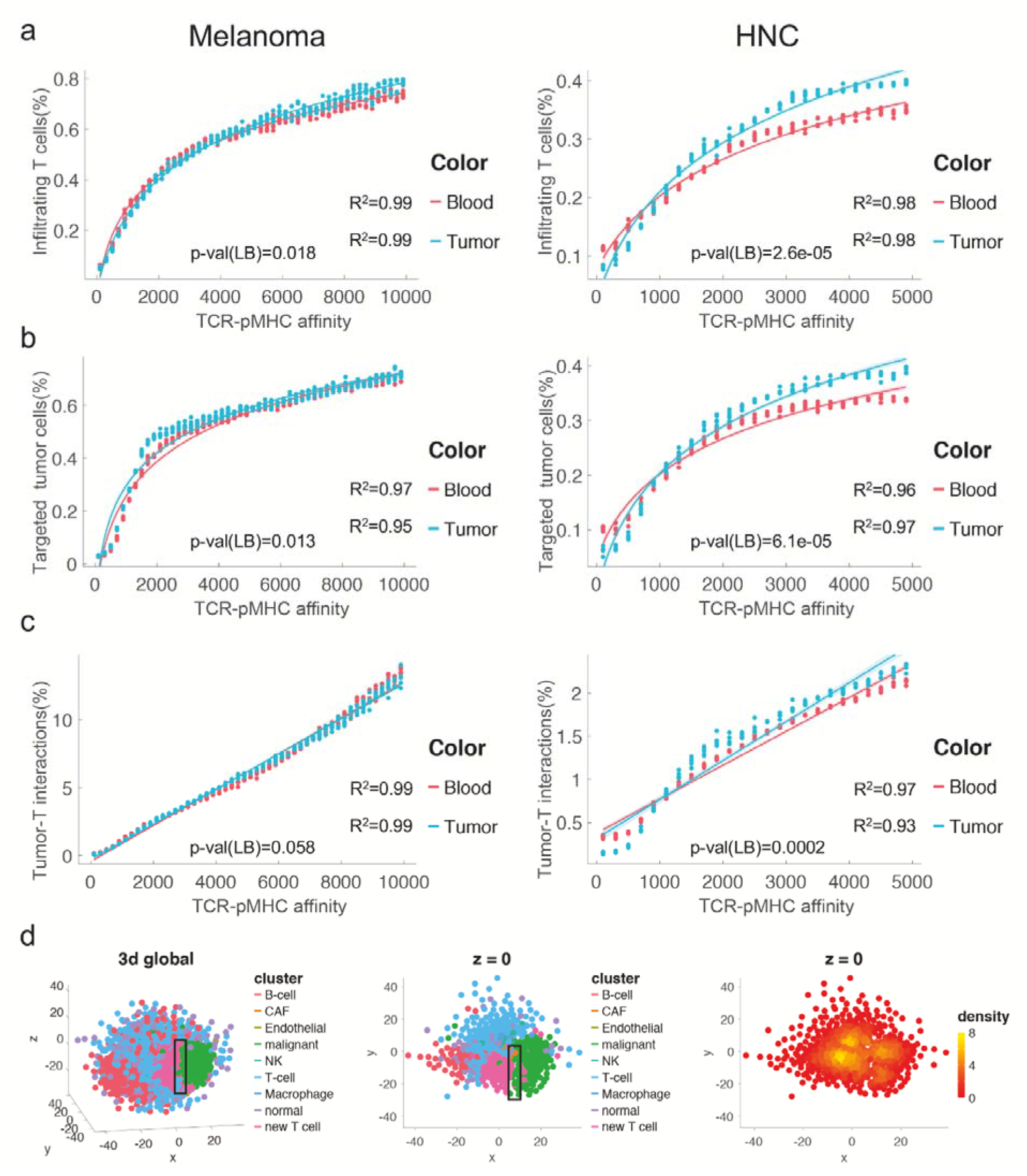
Tumor-T interactions with varying TCR-pMHC affinity after adoptive T cell transfer. (a) The percentage of infiltrating T cells of the melanoma (left) and HNC (right) datasets. “blood” and “tumor” indicate the tissue origins of the adoptively transferred T cells. (b) The percentage of targeted tumor cells of the melanoma (left) and HNC (right) datasets. (c) The percentage of tumor-T interactions relative to the theoretical numbers of the melanoma (left) and HNC (right) datasets. The R^2^ values in (a) and (b) indicated the goodness of fitting a logarithmic function to the observed values of each series. The R^2^ values in (c) indicated the goodness of fitting a linear function to the observed values of each series. The statistical significance of TCR-pMHC affinity and the tissue origins of adoptively transferred T cells to the percentages of infiltrating T cells, targeted tumor cells, and tumor-T interactions was evaluated by ANOVA analysis with repeated measures (by the ranova function of Matlab R2016b), and the lower bound (LB) of the p-values of the tissue origin was displayed. The different dynamics of tumor-T interactions from infiltrating T cells and targeted tumor cells explain the visual patterns of tumor-T interfaces and tumor and T compartments frequently observed in IHC and multiplexed ion beam images (d). New T cell: adoptively transferred T cells.

**Supplementary Fig. 10.**
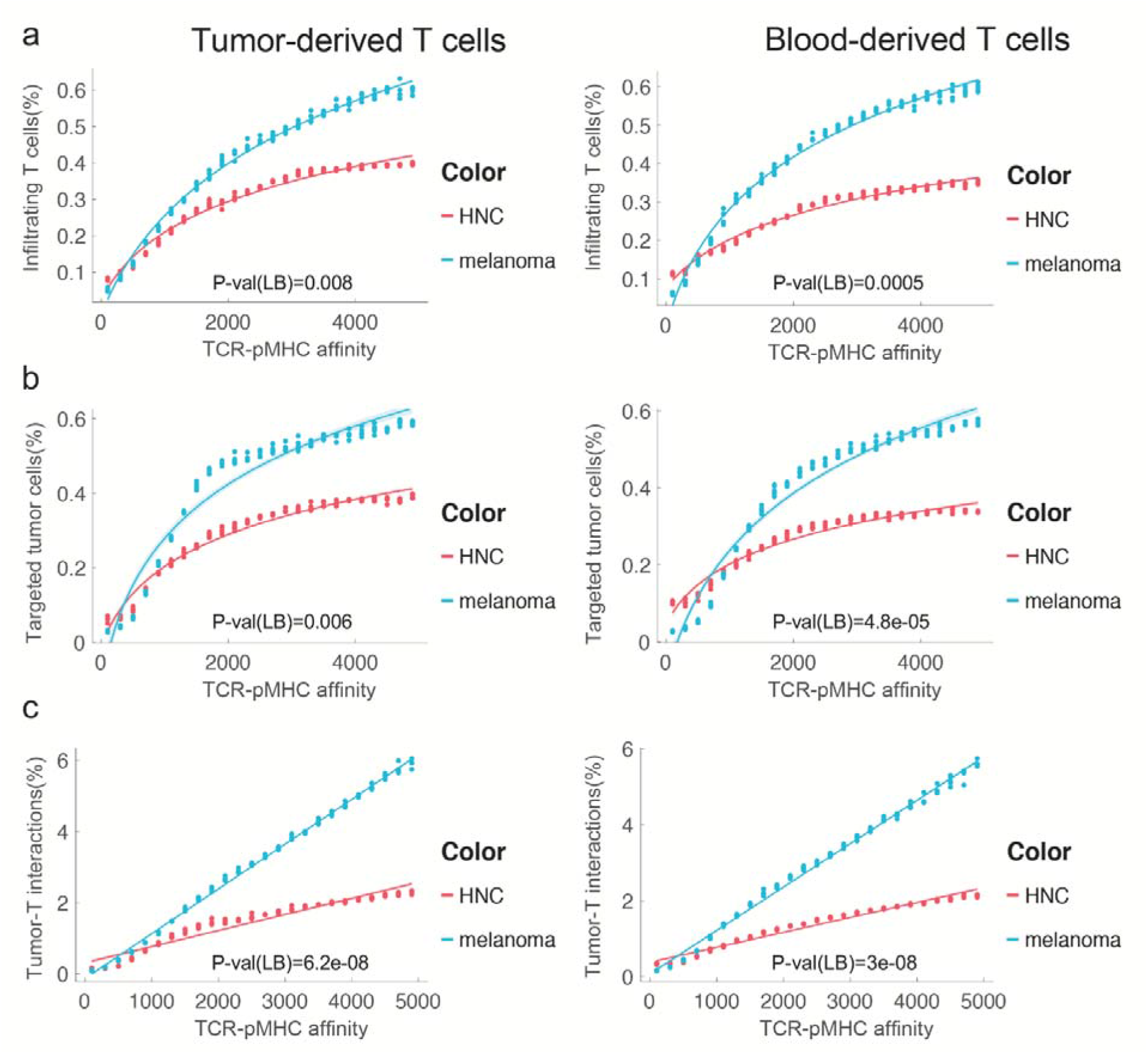
Comparison of the effects of cancer characteristics on tumor-T interactions after adoptive T cell transfer. (a) The percentage of infiltrating T cells of the tumor-(left) and blood-derived (right) T cell transfer. (b) The percentage of targeted tumor cells of the tumor-(left) and blood-derived (right) T cell transfer. (c) The percentage of tumor-T interactions relative to the theoretical numbers of the tumor-(left) and blood-derived (right) T cell transfer. The statistical significance of TCR-pMHC affinity and the cancer characteristics to the percentages of infiltrating T cells, targeted tumor cells, and tumor-T interactions was evaluated by ANOVA analysis with repeated measures (by the ranova function of Matlab R2016b), and the lower bound (LB) of the p-values of the cancer types was displayed.

